# Clinical relevance of zebrafish for gene variants testing. Proof-of-principle with *SMN1/*SMA

**DOI:** 10.1101/2025.01.30.632288

**Authors:** Brett W. Stringer, Yougang Zhang, Afsaneh Taghipour-Sheshdeh, Shuxiang Goh, Heike Kölbel, Michelle A Farrar, Brunhilde Wirth, Jean Giacomotto

## Abstract

Spinal muscular atrophy (SMA) results from *SMN1* gene loss-of-function (LOF), with disease severity directly linked to the level of remaining SMN protein. Nusinersen, risdiplam and onasemnogene abeparvovec are revolutionary treatments but should ideally be implemented before clinical symptoms appear. Because of this, prenatal and newborn screenings are increasingly used to identify common *SMN1* mutations and patients requiring therapy. However, for novel mutations, clinicians lack robust analytic tools to predict pathogenicity before irreversible damage occurs.

To address this gap, we deployed a zebrafish model presenting *smn1*-LOF, exhibiting progressive motor defects and death by only six days of age. We evaluated two *SMN1-*variants (VUS) identified in newborn patients awaiting definite diagnosis and treatment recommendations.

We demonstrated that while known pathogenic variants did not change the disease course, wild-type *SMN1* and both patient variants rescued SMA hallmarks in zebrafish, demonstrating the relevance of this approach for VUS-testing within a crucial timeframe for patients. Both VUS turned out to be non-pathogenic, and therapeutic costs of >US$2 million per child were avoided.

Beyond SMA, this study provides robust proof-of-principle that the zebrafish represents a powerful translational tool for VUS-analysis, and that such approaches should be considered in clinical settings for supporting diagnosis and treatment decisions.

## Introduction

Spinal muscular atrophy (SMA) is a genetic disorder characterized by loss of motor neurons in the spinal cord and brain stem, leading to progressive muscle weakness and atrophy. It is caused by biallelic deletions or pathogenic variants of the survival motor neuron 1 (*SMN1*) gene, which results in a deficiency of the survival motor neuron (SMN) protein, which is crucial for motor neuron function and survival.^1^ SMA severity varies, directly influenced by the *SMN2* gene copy number, which is highly variable among individuals^2^. Without treatment, individuals with the most common form of the disorder, SMA type 1, never gain the ability to sit or stand and usually die or require permanent ventilation within the first 2 years of life (**Fig. 1A**)^1,3-5^.

**Figure 1.**
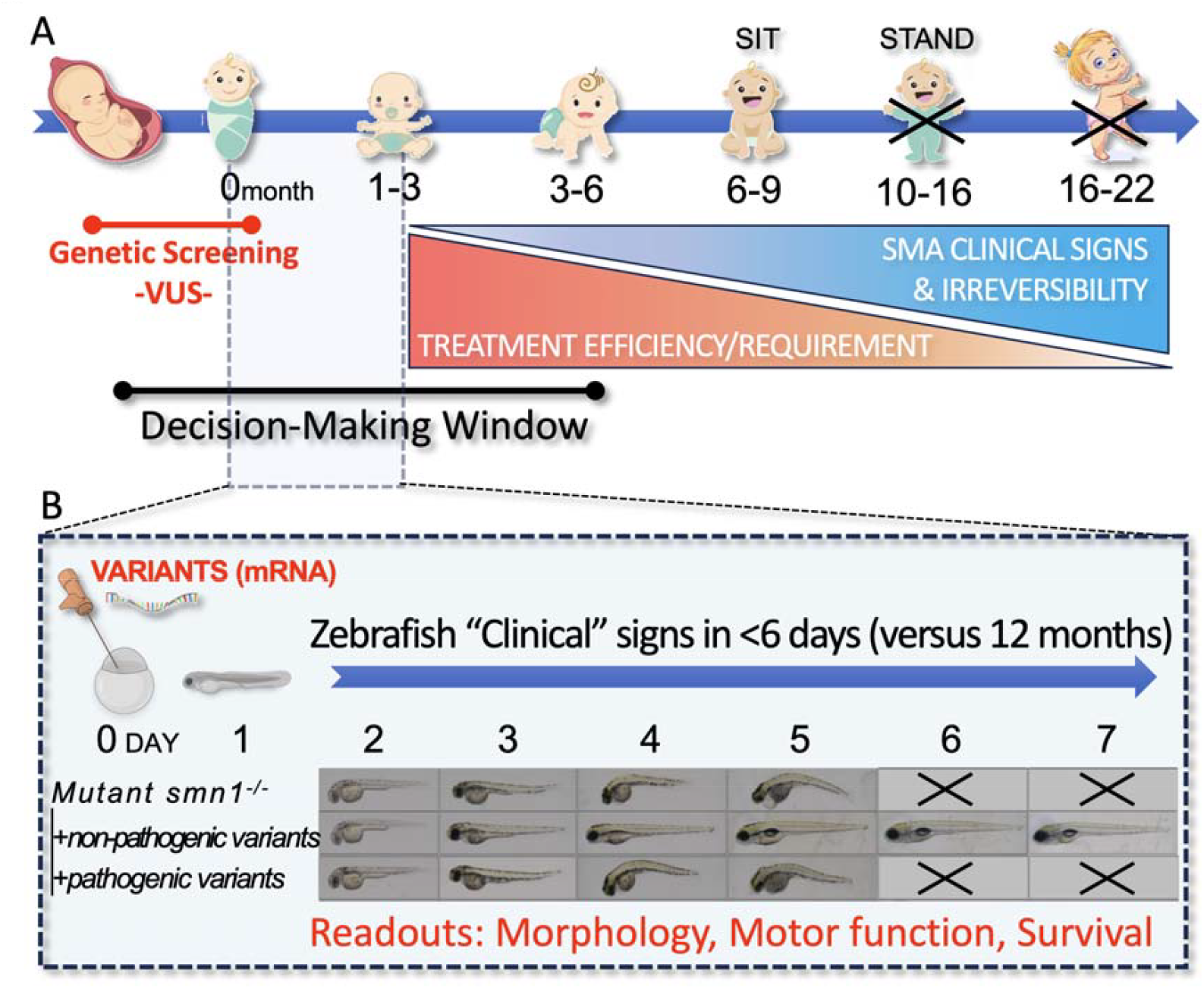
Zebrafish in vivo functional assays can rapidly provide precious information to clinicians regarding the pathogenicity of variants of uncertain significance (VUS). (**A)** Schematic timeline of infant development highlighting the clinical/timeframe dilemma for detection and confirmation of *SMN1*-VUS pathogenicity for early onset SMA (Type I/II), *i*.*e*. treatments need to be implemented before the first major clinical signs. Available treatments are expensive and potentially harmful if unnecessary, emphasizing the need for rapid and innovative VUS-testing approaches within a timeframe that aligns with disease progression. **(B)** Zebrafish functional/complementary assays can quickly provide valuable information on VUS-pathogenicity in less than three months, supporting clinicians in their decision process and in a clinically helpful timeframe. This rapid testing framework is applicable not only to spinal muscular atrophy (SMA) but also to a wide range of pediatric diseases, offering significant benefits in clinical practice.

Until very recently, SMA was the most common inherited cause of infant mortality worldwide.^1^ Thanks to intensive research efforts, three treatments have now been approved and are changing the life of patients and their families. These treatments include the antisense oligonucleotide nusinersen, the small molecule splicing modifier risdiplam, and the gene therapy onasemnogene abeparvovec, all of which aim to increase SMN protein levels.^1,4,5^

However, to be optimally effective, these treatments should be implemented in pre-symptomatic patients, *i*.*e*. prior to the appearance of any major SMA signs or symptoms.^6-9^ Indeed, when patients present with symptoms, irreversible motoneuron death has already occurred (**Fig. 1A**).

To address this dilemma, efforts are being made to diagnose SMA as early as possible and a number of countries have now adopted prenatal and/or newborn screening to identify common pathogenic *SMN1* mutations and patients to be treated. The majority (95%) of individuals with SMA are associated with biallelic *SMN1* exon 7 deletions.^1,4^ Consequently, this is the target analyte in population screening programmes. However, in some cases, novel variants (variants of uncertain significance, VUS) are identified, and the question of their pathogenicity is thereby crucial. Indeed, if pathogenic, definitive confirmation to enable treatment implementation before phenoconversion is important. On the contrary, confirming a variant is non-pathogenic will alleviate psychosocial distress and clinical surveillance.^10^ Unnecessary treatment implementation could also be harmful and significantly expensive; *e*.*g*. onasemnogene abeparvovec costs more than US$2 million for a single dose, and nusinersen more than US$4 million a decade.^1,11,12^

Clinicians and patients would strongly benefit from the development of innovative methodology to help support VUS resolution. Here, we demonstrate that functional/complementation assays using zebrafish are a powerful complementary readout to support clinical decisions for the early-onset SMA forms, and, most importantly, fit within a timeframe that is congruent with pathophysiology (**Fig. 1**).

## Materials and methods

### CASES/PATIENTS

Case 1 was a 12-day-old asymptomatic male infant with no reported family history of SMA, identified through the New South Wales Newborn Screening Programme.^13^ The initial qPCR PerkinElmer NBS assay identified 0 *SMN1* and second-tier ddPCR assays identified 1 *SMN1* and 1 *SMN2* copy. This result was interpreted as an uncertain newborn screening (NBS) for SMA, and the family was contacted for further testing. Diagnostic testing, including *SMN* sequencing, determined that the proband was a compound heterozygote. The infant had a common *SMN1* exon 7 large deletion and an intragenic 4-base pair deletion in exon 7, NM_000344.3, NP_000335.1: c.861_864del (p.Arg288Alafs*5) (**Fig. 2**). The latter mutation was located under the primer used in the first-tier NBS assay, but was not in the region detected by the ddPCR assay. This intragenic *SMN1* mutation was classified as a variant of uncertain significance, VUS-3A, and is referred to as 861VUS in this manuscript.

**Figure 2.**
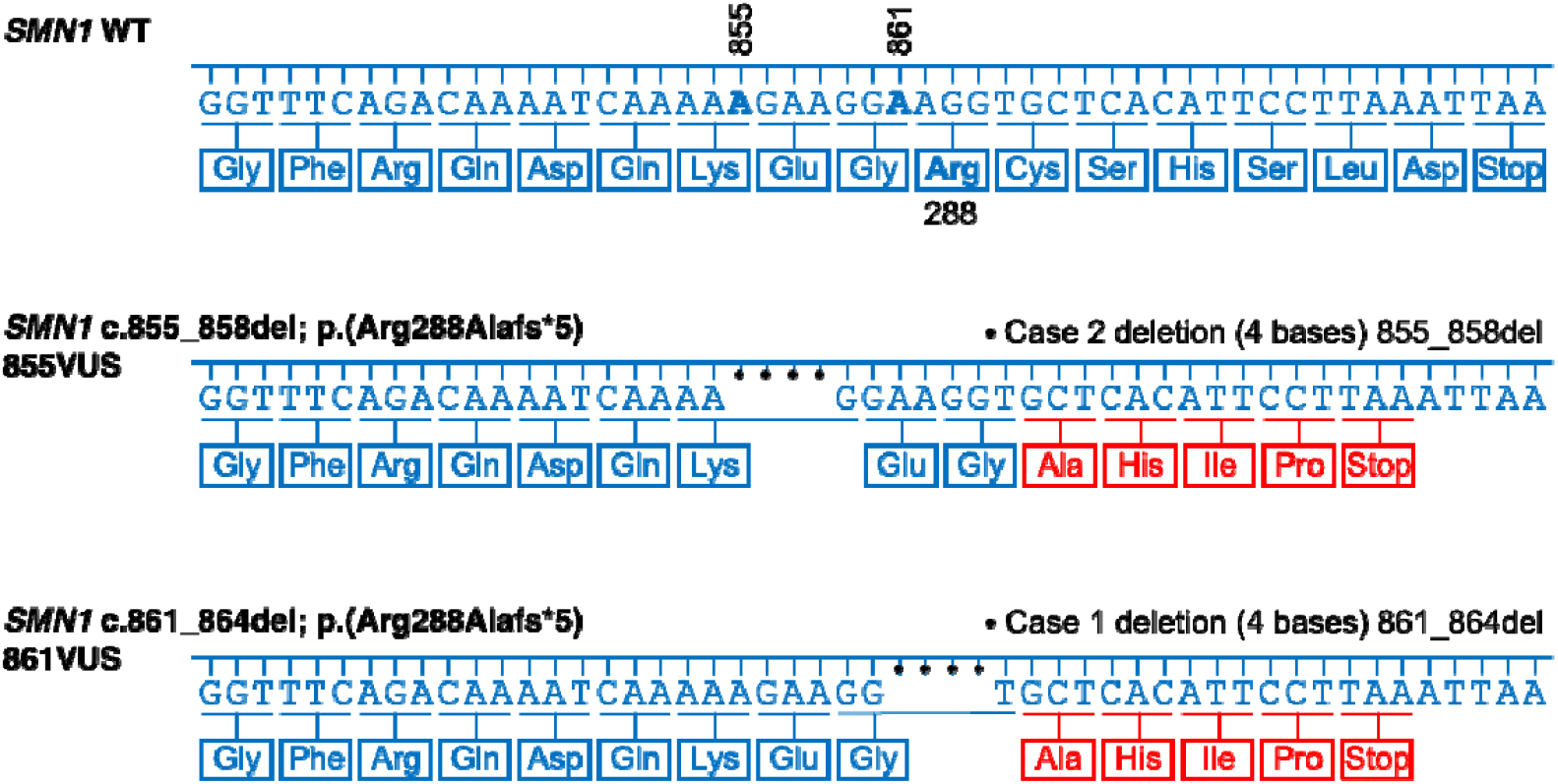
Comparison of wild-type, 855VUS and 861VUS *SMN1* nucleotide and amino acid sequences. The three panels show the nucleotide sequence of the coding strand of the wild-type (WT), 855VUS and 861VUS *SMN1* alleles, from the first nucleotide of the final coding exon to the position of the Stop codon. The 3’ amino acid sequence of the wild-type SMN protein is shown in blue, with changes associated with each VUS in red. The location of coding sequence nucleotides 855 and 861 and amino acid 288 are highlighted in the wild-type strand/sequence and deletions are marked with dots.

Case 2 was a 7-day-old asymptomatic female infant with no reported family history of SMA, identified with 0 *SMN1* via the Newborn Screening Programme in Germany^14^. Routine subsequent genetic validation analysis at the Institute of Human Genetics in Cologne, determined a compound heterozygous *SMN1* genotype with a large *SMN1* and *SMN2* deletion on one haplotype and an intragenic 4-base pair deletion in exon 7, *SMN1* ENST00000380707 c.855_858del p.(Arg288Alafs*5) and absence of any *SMN2* copy on the second haplotype (**Fig. 2**). The intragenic *SMN1* mutation was classified as a variant of uncertain significance, VUS-3A. This VUS is named 855VUS in this manuscript.

Interestingly, although independent and different, both *SMN1* variants are predicted to encode the same truncated protein, with the last seven amino acids truncated and substituted with four different amino acids before a premature stop (**Fig. 2**).

### ZEBRAFISH

Zebrafish were maintained by standard protocols approved by Griffith University, approval GRIDD1122AEC. Homozygous *smn*^*-/-*^ zebrafish embryos were produced as described previously.^15,16^

### GENETIC AND FUNCTIONAL ANALYSIS

*SMN1* wild-type and variant cDNAs were synthesized as gene blocks by Gene Universal *Inc*. and inserted into a pME Gateway-compatible backbone (**Supplementary Table 1**). Each pME-*SMN1* was further subcloned into a custom RNA expression plasmid (Addgene #171793)^17^ by Gateway cloning. mRNAs were synthesized using a mMESSAGE mMACHINE transcription kit (Invitrogen) from plasmid DNA linearized by *Kpn*I digestion, purified with a MEGAclear™ Clean-Up Kit (Invitrogen) according to manufacturer protocols, and stored at -80°C. Purified mRNAs were microinjected into the yolk of 1-cell stage embryos at 250 pg final as previously described.^18^ Zebrafish morphology was recorded using an Olympus MVX10 microscope driven by cellSens software. Survival was recorded each morning at 9am, death was determined by absence of a heartbeat. To assess motor function, larvae were distributed in 24-well plates with a single animal per well and supplemented with 0.5 mL of E3 medium. Motor functions were assessed for 24 minutes using a ZebraBox Revolution (ViewPoint). The protocol included repetitive cycles of 4 minutes of light and 4 minutes of dark (3 cycles). The protocol was complemented with repetitive flashes of light and 1-second acoustic vibration (250 Hz) every minute (independently offset by 30 seconds) to assess swimming reactions/abilities.

## Data availability

Data presented in this study are available upon request.

## Results

Two cases, respectively carriers of *SMN1*-VUS c.861_864del (**861VUS**) and c.855_858del (**855VUS**) (**Fig. 2**) were identified at 2 weeks-of-age and brought to our attention by the Australian Functional Genomics Network (AFGN). Each case/patient was simultaneously a carrier of a common pathogenic deletion of *SMN1* exon 7, putting them at risk of developing SMA. Moreover, one newborn failed to carry any *SMN2* copy, the other carried one *SMN2* copy, suggesting the potential to develop a severe SMA type. However, no clinical symptoms were present at the time of discussion, and determination of the possible pathogenicity of each VUS was important to support either a decision to implement treatment or to possibly delay it until the emergence of the first subclinical electrophysiological SMA hallmarks/diagnosis. To address the potential pathogenicity of these VUS, we deployed a zebrafish SMA model deficient in SMN function and tested independently whether each variant could successfully complement the lack of SMN function and associated phenotypes. (**Fig. 1**).

### Patients’ VUS rescued morphological defects of SMA zebrafish

To determine whether the *SMN1* variants were pathogenic, we investigated whether *in vitro*-transcribed mRNAs from 861VUS and 855VUS, injected into single-cell *smn*^-/-^ zebrafish embryos, successfully rescued the previously described zebrafish SMA hallmarks, including morphology, swimming ability, and survival (**Fig. 3 and 4**).^16,18,19^ To evaluate the efficiency of our approach, we used human wild-type *SMN1* mRNA (“WT” – positive control) that we previously demonstrated to be efficient in rescuing zebrafish SMA phenotypes^16,18,19^. For robustness, we included a series of additional controls, including (i) mock-injected *smn*^-/-^ (“*smn*^*-/-*^” - negative control), (ii) stable transgenics expressing human wt-*SMN1* mRNA ubiquitously (“TgSMN1” - positive control)^16^, (iii) known pathogenic human *SMN1* mRNA variant (*SMN1* c.549del) (“path” – negative control) and (iv) non-pathogenic human *SMN1* mRNA variant (*SMN1* c.462A>G) (“non-path” - positive control).

**Figure 3.**
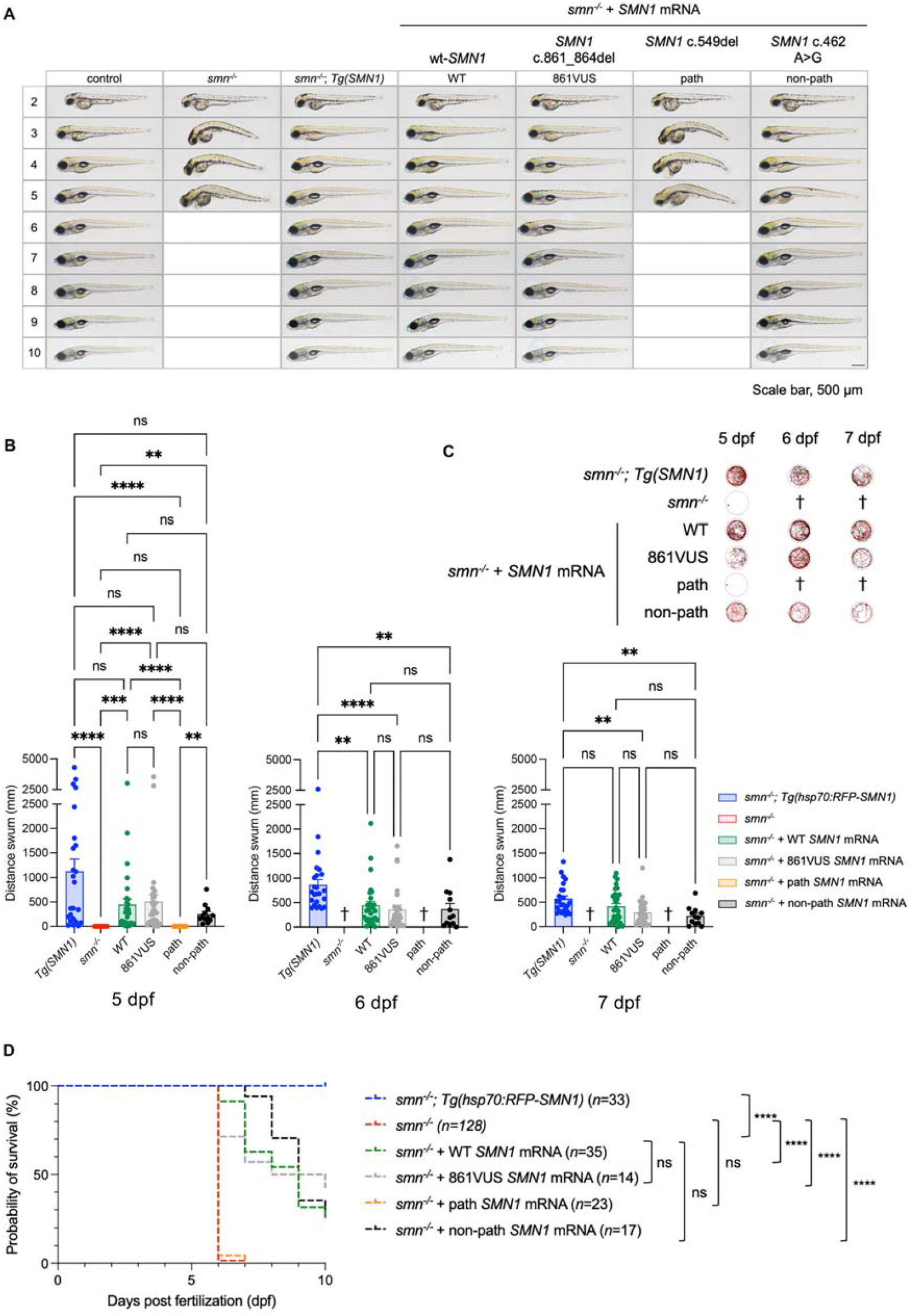
861VUS mRNA injections rescue SMA *smn*^-/-^ zebrafish disease hallmarks. **(A)** *smn*^-/-^ zebrafish develop morphological malformations from 3 dpf, being smaller with a progressive body curvature and small eyes. From 4 dpf, animals develop progressive pericardial and cerebral oedema. Animals died between 6 and 7 dpf, as determined by the absence of a heartbeat. mRNA from the known pathogenic *SMN1* variant, *SMN1* c.549del (path – negative control), as expected, failed to rescue this phenotype (did not prevent the appearance of these malformations). In contrast, positive controls, (i) ubiquitous transgenic expression of wt-*SMN1* (*Tg(SMN1)*), (ii) injected wt-*SMN1* mRNA (WT), (iii) injected non-pathogenic variant c.462A>G (non-path), and importantly (iv) injected c.861_864del *SMN1* mRNA (861VUS) all rescued these morphological traits, with no distinguishable difference between the animals. These results robustly demonstrate that 861VUS produces a functional SMN protein. Empty boxes indicate 100% batch mortality. Scale bar, 100 µm. **(B)** SMA *smn*^*-*/-^ zebrafish present dramatic motor function loss at 5 dpf and die between 6 and 7 dpf. As previously demonstrated, human wt*-SMN1* transgenic ubiquitous expression (*Tg(SMN1)*) or injected wt-*SMN1* mRNA (WT) rescue this motor function loss. Similarly, both the non-pathogenic *SMN1* variant (non-path) and 861VUS efficiently restore motor function with no detectable significant difference. On the contrary, confirming the robustness of the approach, mRNA from the pathogenic *SMN1* variant (path) failed to improve motor function with no difference from negative control (*smn*^-/-^). Graphs represent comparisons of the total distance swum by each cohort of fish over 24 minutes. Each graph summarizes three independent biological replicates. n=12-30 fish per condition. Kruskal-Wallace test with Dunn’s correction for multiple comparisons. *\*\*\*\*, P*<0.0001; *\*\*\*, P*<0.001; *\*\*, P*<0.01; *\*, P*<0.05; ns, not significant. **(C)** Representative swimming tracks at 5, 6 and 7 dpf. Video samples are available in supplementary data. Cross symbols indicate 100% mortality. **(D)** SMA *smn*^-/-^ zebrafish (*smn*^-/-^) had a median survival of 6 days. Injected mRNA from the known pathogenic *SMN1* variant (path) did not have any effect on survival. In contrast, demonstrating restoration of Smn/SMN function, injected mRNA from 861VUS extended survival similarly to wt-mRNA (WT) and non-pathogenic variant (non-path) animals; all extended the survival of *smn*^*-/-*^ larvae. *\*\*\*\*, P*<0.0001; ns, not significant. Log-rank (Mantel-Cox) test.

**Figure 4.**
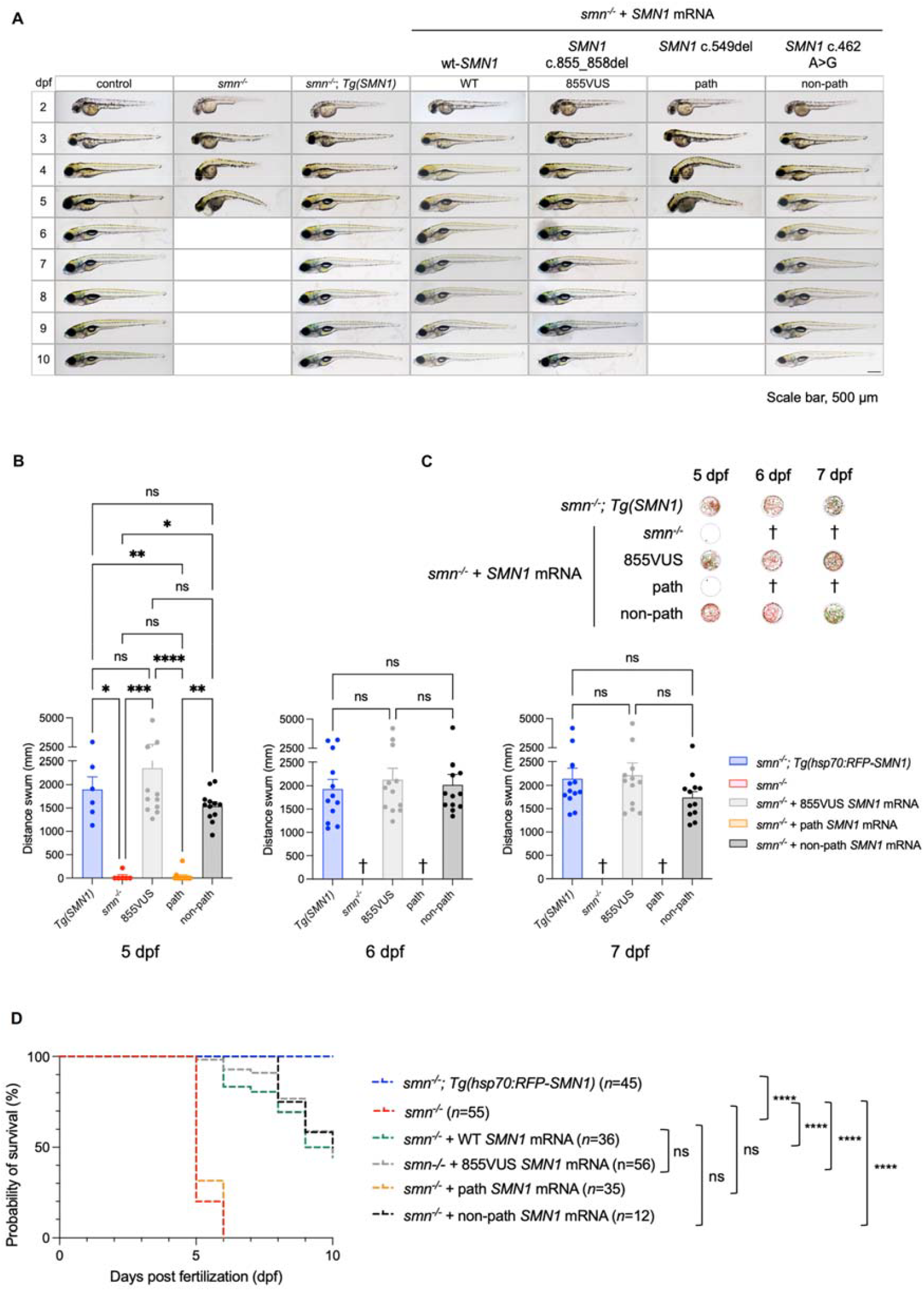
855VUS mRNA injections rescue SMA *smn*^-/-^ zebrafish disease hallmarks. **(A)** Morphological malformations consistent with spinal motor neuron degeneration and skeletal muscular atrophy were again evident in *smn*^-/-^ embryos, which from 3 dpf were smaller with a curved body and small eyes. Pericardial and/or cerebral oedema appeared at 4 and 5 dpf. mRNA from the known pathogenic *SMN1* variant, *SMN1* c.549del (path), again as expected failed to rescue this phenotype. In contrast, injected mRNA from the *SMN1* variant of uncertain significance, *SMN1* c.855_858del, (855VUS), as well as injected wt-*SMN1* mRNA (WT) or wt-*SMN1* mRNA from an endogenous *SMN1* transgene (*Tg(SMN1)*), and mRNA from the known non-pathogenic *SMN1* variant *SMN1* c.462A>G (non-path) all rescued the phenotype of *smn*^-/-^ embryos and larvae, providing evidence of functional SMN protein. Empty boxes indicate all larvae are dead. Scale bar, 100 µm. **(B)** *SMN1* c.855_858 mRNA (855VUS) rescued the swimming ability of *smn*^*-*/-^ larvae (*smn*^*-*/-^) just as well as wt-*SMN1* mRNA expressed from an endogenous transgene (*Tg(SMN1)*), and mRNA from the non-pathogenic *SMN1* variant c.462A>G (non-path). mRNA from the pathogenic *SMN1* variant, *SMN1* c.549del (path), in contrast did not. Comparison of the total distance swum by each cohort of fish over 24-minutes. Each graph summarizes three independent biological replicates. n=12 fish per condition. Kruskal-Wallace test with Dunn’s correction for multiple comparisons. *\*\*\*\*, P*<0.0001; *\*\*\*, P*<0.001; *\*\*, P*<0.01; *\*, P*<0.05; ns, not significant. **(C)** Representative examples at 5, 6 and 7 dpf of the swimming tracks/patterns for each cohort of fish over 24 minutes. Individual movement tracks begin at the orange square and end at the black square. Video examples shown in supplementary data. Cross symbol indicates no surviving fish. **(D)** *smn*^-/-^ larvae (*smn*^-/-^) had a median survival of 6 days. mRNA from the known pathogenic *SMN1* variant, *SMN1* c.549del (path) again did not alter this. In contrast, demonstrating restoration of Smn/SMN function, injected mRNA from the *SMN1* variant of uncertain significance *SMN1* c.855_858del, (855VUS) extended the survival of *smn*^*-/-*^ larvae just as well as injected wt-*SMN1* mRNA (WT) and mRNA from the known non-pathogenic *SMN1* variant *SMN1* c.462A>G (non-path). *\*\*\*\*, P*<0.0001; ns, not significant. Log-rank (Mantel-Cox) test.

As previously shown, and replicated here, absence of Smn/SMN protein in the *smn1*^***-/-***^ SMA zebrafish model triggered morphological malformations from 3 days post fertilization (dpf), with larvae being smaller in size and presenting body curvature and small eyes (**Fig. 3A**). These hallmarks became more prominent at 4 and 5 dpf. Pericardial and cerebral oedema appeared from 4 dpf. Finally, 100% of larvae died between 6 and 7 dpf, as identified by the absence of heartbeats. None of the negative controls, mock-injection and pathogenic variant injections (path) modified these disease hallmarks, with all embryos and larvae exhibiting the same malformations and with similar dynamics/progression (**Fig. 3 and 4**). In contrast, and confirming the robustness and relevance of our methodology, mRNAs from both wt-*SMN1* (WT) and a non-pathogenic *SMN1* variant (non-path) rescued these phenotypes. Importantly, mRNAs from both 861VUS and 855VUS also rescued these hallmarks, with all embryos and larvae having indistinguishable development/morphology when compared to all positive controls (WT, TgSMN1, non-path). This was clearly apparent from 3 dpf and persisted to 10 dpf, when the experiment was ended.

Together, these data demonstrate that mRNA from both *SMN1* VUS rescued the morphological phenotype of *smn*^-/-^ SMA zebrafish as effectively as wild-type *SMN1*, suggesting that each VUS properly translates into a functional SMN protein.

### Patients’ VUS improved motor functions of SMA zebrafish as efficiently as wt-*SMN1*

To further investigate the function/pathogenicity of the identified VUS, we conducted functional assays assessing motor function recovery following SMN complementation. Swimming behavior and ability of zebrafish larvae were analyzed using automatic tracking at 5, 6, and 7 dpf as per the method section. At 5 dpf, mock-injected SMA *smn*^*-/-*^ larvae (*smn*^-/-^) exhibited dramatic motor function loss, as demonstrated by a near-absent spontaneous swimming pattern and response to stimulation (**Fig. 3B/C and 4B/C)**. As expected, injections of the pathogenic variant (path) failed to rescue these motor function losses and were not significantly different from mock-injected SMA *smn*^*-/-*^ control animals. In contrast, injection of either wt-*SMN1* (WT), non-pathogenic variant (non-path), and both patients’ 861VUS and 855VUS, robustly restored motor function as demonstrated by significantly longer distance swum (**Fig. 3B/C and 4B/C**).

It is noteworthy that at 6 dpf, all negative control larvae were dead (mock-injection and pathogenic variant), demonstrating proper progression of the disease and execution of the experiments. In contrast, *smn*^*-/-*^ larvae in the remaining cohorts remained alive and healthy, indistinguishable from injected positive controls, and continued to swim without statistically significant difference at both 6 dpf and 7 dpf (**Fig. 3B/C and 4B/C**).

Together, these data demonstrate that both patients’ VUS mRNAs rescue motor function of SMA *smn*^-/-^ zebrafish as efficiently as wt-*SMN1* mRNA or a non-pathogenic *SMN1* variant.

### Patients’ VUS increased survival of SMA zebrafish as efficiently as wt-*SMN1*

We complemented our readout/assays by investigating the effect of both 861VUS and 855VUS on survival. In the absence of Smn/SMN protein, SMA *smn*^*-/-*^ zebrafish (*smn*-/-) have a highly replicable median survival of 6 days-post-fertilization (**Fig. 3D and 4D**).^16^ While the addition/injection of pathogenic *SMN1* variant mRNA (path) failed to modify this mortality in all experiments conducted, the wt-*SMN1* (WT), non-pathogenic *SMN1* (non-path) and patients’ VUS (861VUS and 855VUS) extended the median survival to over 9 days. These data robustly support that the patients’ VUS retain biological function and do not differ statistically in our assay from the positive controls based on wt-*SMN1* and a non-pathogenic *SMN1* variant. Experiments were stopped at 10 dpf as injected mRNAs have a time-restricted efficacy, being degraded in the first week after injection.^20,21^

Taken together, these data show that both patients’ VUS rescued all tested SMA hallmarks in the zebrafish (morphology, motor function, and survival), clearly indicating that the protein encoded by these novel variants retains significant biological function.

Fully confirming the relevance of the presented methodology, the patients were around six months of age at the time of experiment completion and were still asymptomatic. After review and consultation with all other evidence, a decision was made to not implement treatment and wait until the patients were twelve months old to write this manuscript, at which time both infants remained asymptomatic, exhibiting normal motor, neurological and electrophysiology examination. Both patients achieved all motor milestones at expected timepoints, with patient 1 attaining independent walking at 14 months of age.

## Discussion

In recent years, the introduction of nusinersen, risdiplam, and onasemnogene abeparvovec has dramatically transformed the landscape of SMA, which was one of the most common inherited causes of infant mortality worldwide. For early-onset/severe forms of SMA, these treatments are most effective when administered before the onset of clinical symptoms (**Fig 1**).^1,22^ This critical time window demands early and accurate diagnosis of SMA, which has been facilitated by widespread prenatal and newborn screening programmes for the detection of common/known mutations. However, the interpretation of novel gene variants of uncertain significance (VUS) remains a formidable challenge in the clinical setting. Our study demonstrates the potential of zebrafish as a powerful translational model for the functional characterization of *SMN1* VUS, providing essential data to support both diagnosis and clinical decision-making.

We investigated two *SMN1* VUS identified in neonates through newborn screening. These two different variants, c.861_864del (861VUS) and c.855_858del (855VUS), are predicted to encode the same truncated SMN protein. Our functional assays demonstrated that, when injected in the early embryo, mRNAs from 861VUS and 855VUS could efficiently rescue the morphological, motor function, and survival defects in SMA *smn*^*-/-*^ zebrafish larvae as effectively as human wild-type *SMN1* mRNA. These findings suggest that the proteins encoded by these variants retain sufficient biological function to compensate for the loss of endogenous SMN protein, thus providing crucial evidence for their non-pathogenicity. Because both VUS proved to be non-pathogenic, the initiation of drug therapy with potential side effects and the physical and psychological stress for the child and parents could be avoided. In addition, the public health system did not bear the therapy costs of around US$2 million (or more) per child.

As a guide, the assessment of these *SMN1* variants was completed in 18 working days (**Fig. 1**). This did not include the initial synthesis of coding DNA sequence (cds) by an external supplier, a time constraint that in future should add no more than approximately a week to the pipeline. Functional evaluation of *SMN1* RNA in *smn*^-/-^ zebrafish embryos and larvae was the shortest step of the process, and clear evidence of SMN function was apparent in as little as six days. In conclusion, this study demonstrates that zebrafish is a robust and inexpensive tool to assess *SMN1*-VUS and should be considered in clinical settings to support diagnosis and the need for treatment.

Importantly, as newborn screening (NBS) for SMA is expanding, additional babies are anticipated to be identified with novel variants. This will cause intense distress for families, rapid and accurate VUS resolution is therefore highly impactful and will help guiding a definitive clinical pathway, either treatment or reduction of surveillance, stress and uncertainty for families and health services.

### Implications for clinical practice and application to other diseases

Going beyond SMA, our study provides robust proof-of-principle that zebrafish can serve as a powerful tool for VUS resolution. To the best of our knowledge, this represents the first demonstration of their successful translational use within a clinically meaningful timeframe. Indeed, our study highlights the clinical relevance of zebrafish as a valuable model for gene variants (VUS) testing, not only for SMA, but for human disease in general. With exome and whole genome sequencing becoming more commonplace, we are seeing an explosion of new gene variants associated with a diversity of disorders. The determination of their functional consequences is posing a significant challenge, and this problem will continue to grow with the development and need for precision/personalised medicine. The development of robust tools for VUS resolution is becoming critical. Our work shows that the zebrafish can help address this translational need.

## Acknowledgements

We thank the children investigated, their families, and carers for their valuable contributions to data collection. We thank the clinical geneticist (Dr. Mert Karakaya, Univ. of Cologne) and the investigators of the genetic diagnoses (Dr. Nico Fuhrmann, Dr. Jutta Becker from Univ. of Cologne) involved in the routine diagnostic investigation of the newborns. We also thank the Australian Functional Genomics Network, which greatly facilitated the initiation of this study.

