## Supplemental table 1 for "Clinical relevance of zebrafish for gene variants testing. Proof-of-principle with *SMN1/*SMA"

Supplementary Table 1: Plasmids created for *SMN1* VUS analysis.

| **Plasmid name** | **Purpose** |
|---|---|
| 249-pME-hsa_SMN1-wt | contains *SMN1* wild-type cDNA |
| 250-pME-c.861_864_SMN1 | contains *SMN1* variant cDNA (861VUS) |
| 252-pME-c.855_858_SMN1 | contains *SMN1* variant cDNA (855VUS) |
| 253-pME-c.549del_SMN1 | contains *SMN1* variant cDNA (path) |
| 254-pME-c.462A>G_SMN1 | contains *SMN1* variant cDNA (non-path) |
| 260-pT3-hsa_SMN1-WT_polyA | in vitro transcription of *SMN1* wild-type mRNA |
| 261-pT3-c.861_864_hsa_SMN1_polyA | in vitro transcription of *SMN1* 861VUS mRNA |
| 263-pT3-c.855_858_hsa_SMN1_polyA | in vitro transcription of *SMN1* 855VUS mRNA |
| 264-pT3-c.549del_hsa_SMN1_polyA | in vitro transcription of *SMN1* path mRNA |
| 265-T3-c.462A>G_hsa_SMN1_polyA | in vitro transcription of *SMN1* non-path mRNA |
